# Aiolos is specifically altered in schizophrenia, is expressed in hippocampal neurons and regulates structural synaptic plasticity through complement-mediated mechanisms

**DOI:** 10.64898/2026.11.22.768424

**Authors:** Maoyu Wang, Marcos Galán-Ganga, Nerea Gómez-Rivada, Laura López-Molina, Pol García-Segura, Laura Martín-Herrero, Cristina Malagelada, Kentaro Yomogida, J Javier Meana, Luis F Callado, Miguel Montero-Escobedo, Myriam Cavero, Carla Hernandez, Ana Escanilla, Belen Ramos, Jordi Alberch, Belén Arranz, Marina López-Solà, Iván Ballasch, Albert Giralt

## Abstract

Aiolos (Ikzf3), a member of the Ikzf family, has been primarily characterized in the immune system, whereas other Ikzf family members have reported roles in the brain and have been implicated in schizophrenia. However, the potential role of Aiolos in the brain remains broadly unexplored. Combining human samples, translational and gain and loss of function mouse models we characterized Aiolos expression in the brain and investigated its potential involvement in schizophrenia. Here we show that Aiolos expression varies across brain regions and developmental stages, stabilizing in principal neurons during adulthood. Moreover, we identified that Aiolos is dysregulated in peripheral immune cells and in the hippocampus of patients with schizophrenia. Modeling the impact of those dysregulations in mice induced behavioral phenotypes relevant to the positive, negative and cognitive symptoms of schizophrenia. Finally, we found that mimicking Aiolos hippocampal dysregulation in mice generated increased complement protein abundance, accompanied by microglia-associated loss of synaptic material. Our results suggest that Aiolos may regulate complement-mediated synaptic elimination with potential implications for schizophrenia.

## INTRODUCTION

Aiolos (Ikzf3) is a member of the Ikaros zing finger protein (Ikzf) family of transcription factors. The family comprises four additional members: Ikaros (Ikzf1), Helios (Ikzf2), Eos (Ikzf4) and Pegasus (Ikzf5) [1]. The functions of Aiolos as well as other members of the Ikzf family (e.i. Ikaros and Helios), have been largely described in the immune context [2], [3], [4]. These transcription factors are essential for the normal development of immune cells, including B, T, and Natural Killer cells. Moreover, they play key roles in regulating cytokines expression such as Il-2, Il-4, Il-12, and IFN-γ among others[5]. Within the nervous system, Aiolos’ siblings Ikaros and Helios can be expressed during neurodevelopment in striatal and cortical neurons [6], [7]. Furthermore, in recent years Ikaros has also been shown to be present in microglia of the adult mouse brain [8]. Dysregulations of these transcription factors have been linked to neuroinflammation, neurodegeneration and psychiatric conditions such as schizophrenia [8], [9], [10]. In contrast, the role of Aiolos in the brain remains broadly unexplored, and its potential involvement in psychiatric conditions is still unknown.

Schizophrenia is a psychiatric condition that impacts various aspects of an individual’s cognition, perception, and behavior [11]. Its prevalence round around 23 million people worldwide and accounts for an important health care burden with an average of 16.500€ of annual associated costs per patient in Europe [12], [13]. Schizophrenia symptoms are classically organized in three broad categories. Positive symptoms, such as hallucination, delusions and thought disorder. Negative symptoms, including social withdrawal and lack of motivation, and cognitive symptoms, comprising deficiencies in executive functions, working memory and processing speed [11], [14]. Schizophrenia pathophysiology is increasingly understood as involving the interplay between neural and immune dysregulations [15], [16], [17]. For example, genetic wide association studies (GWAS) pointed out the existence of polymorphisms in proteins of the complement system as well as altered levels in circulating serum [18], [19], suggesting a potential role for these proteins in processes such as synaptic pruning [20]. On the other hand, research during last decade indicates the presence of persistent neuroinflammation processes in schizophrenia including impairments in glial cells functions [21], [22]. However, the precise mechanisms that underpin these dysregulations and how they contribute to the different kinds of symptoms are still not fully understood. In this context, given its implication in the immune system and the involvement of Ikaros and Helios in schizophrenia, Aiolos arises as a plausible yet largely unexplored contributor to schizophrenia pathophysiology.

In the present work, we characterized Aiolos across development in different brain regions and found that its levels rise during perinatal stages and become stable in youth-adulthood. Moreover, we identified that in adulthood Aiolos is mostly expressed in mature principal neurons. Using loss-of-function and gain-of-function strategies we observed that Aiolos is a key regulator of several complement proteins, suggesting potential implications for schizophrenia. Indeed, here we show that Aiolos levels are altered in both peripheral blood mononuclear cells (PBMCs) and in hippocampal postmortem samples of patients with schizophrenia. Finally, manipulating Aiolos levels in mouse hippocampi we induced behavioral phenotypes recapitulating positive and cognitive schizophrenia-like symptomatology. Our findings further suggest Aiolos-driven complement implication in microglia-mediated loss of synaptic material.

## METHODS

### Human post-mortem brain samples

The brain samples from schizophrenia (SCZ) patients used in this study were provided by the Sant Joan de Déu Brain Bank (Sant Boi de Llobregat, Barcelona, Spain). The donation and obtention of samples were regulated by the ethics committee of the institution. The sample processing followed the rules of the European Consortium of Nervous Tissues: BrainNet Europe II (BNEII). All the samples were protected in terms of individual donor identification following the BNEII laws. Clinical diagnosis of donor subjects was confirmed pre-mortem with DMS-IV (Diagnostic and Statistical Manual of Mental Disorders, 4th edition) and ICD-10 (International Statistical Classification of Diseases and Related Health Problems) criteria by clinical examiners. Control samples were obtained from Banc de Teixits Neurològics (Servei Científico-Tècnics, Universitat de Barcelona, Barcelona, Spain) and the sample processing also followed the BNEII rules. Post-mortem brain tissue samples from individuals with MDD and age-matched controls, all with a post-mortem delay (PMD) below 24 h, were obtained at autopsy in the Basque Institute of Legal Medicine in Bilbao, Spain. Samples from MDD subjects and age-matched controls were collected from the hippocampus. Samples were immediately stored at −80 °C. Medical records were reviewed retrospectively to identify individuals with MDD diagnoses. All diagnoses were made antemortem by certified psychiatrists of the Basque Healthcare System (Osakidetza) following Diagnostic and Statistical Manual of Mental Disorders, Fourth and Fifth Editions (DSM-IV and DSM-5), or the International Classification of Diseases, Tenth Revision (ICD-10) criteria. All the procedures for the obtention of post-mortem samples followed the ethical guidelines of the Declaration of Helsinki and local ethical committees (SCH: Universitat de Barcelona: IRB00003099; Fundació CEIC Sant Joan de Déu: BTN-PSSJD, CER122306 and MDD: University of Barcelona: Institutional Review Board IRB00003099, CER122306). Demographic information can be found in supplementary Tables 1 and 2.

### Recruitment of patients

The blood samples from schizophrenia patients were obtained in the Outpatient clinic (N◦PI17/00246, PI Belen Arranz) located in Cornellà, Barcelona, Spain (Parc Sanitari Sant Joan de Deu). Adult controls and patients with DSM-5 (Diagnostic and Statistical Manual of Mental Disorders) schizophrenia-spectrum disorder at any stage of the disease were included. The inclusion criteria for patients were: (1) adults over 18 years of age; (2) ability to speak Spanish correctly; and (3) signed informed consent. Exclusion criteria were: (1) history of head trauma with loss of consciousness and (2) organic disease with mental repercussions; (3) presence of an acute inflammatory process: (3.1) fever (> 38◦C) or infection in the 2 weeks prior to the baseline interview, or (3.2) have received vaccinations in the past 4 weeks. This study was conducted following the ethical principles of the Declaration of Helsinki and Good Clinical Practice and the Ethics and Research Board from Parc Sanitari Sant Joan de Deu. All participants provided written informed consent prior to their inclusion in the study. Demographic information can be found in supplementary Table 3.

Regarding the mayor depressive disorder (MDD) patients, all participants were female and met a diagnosis of MDD or adjustment disorder subtype depressive / mixed, following the Diagnostic and Statistical Manual of Mental Disorders - 5th Edition criteria for a current depressive episode assessed by a licensed psychiatrist, as well as a score in the Patient Health Questionnaire-9 (PHQ-9) ≥9 [23], indicating at least moderate depression. Healthy controls had no current or previous diagnosis of psychiatric disorders and scored below the clinical cut-offs on the main MDD-related questionnaires (PHQ-9 < 9; FIQ < 39). Demographic information can be found in supplementary Table 4.

### Human peripheral blood mononuclear cells (PBMCs)

Human peripheral blood mononuclear cells (PBMCs) were isolated from peripheral blood. For this procedure, 10 ml blood samples were diluted with 10 ml of phosphate-buffered saline (PBS) pH-7.2. Consequently, 10 ml of diluted blood were placed in 15 ml tubes filled with 4 ml of a density gradient medium with ρ = 1.077 g/ml (e.g. Ficoll-Paque PLUS) and centrifuged at 500 × g for 35 min at 20 °C without brake. PBMCs were then transferred to 50 ml tubes, the tubes were filled with PBS and centrifuged at 500 × g for 10 min at 20 °C. PBMCs were then lysated for mRNA (see next section) or plated. For PBMCs plating, after elimination of the supernatant, PBMCs were resuspended in X-vivo medium (#BEBP02-055Q Lonza bioscience, Maryland, USA) supplemented with pen/ strep 1%/1%, l-glutamine 1% and Hepes 0,02 M. Cells were seeded in 24-well plates (1 ml per well at 4·10⁶ cells/ml) and left in an incubator at 37 °C, 5% CO2. After 24 h, PBMCs were treated with PMA (50 ng/ml, #P1585, Sigma-Aldrich Chemical Co., St. Louis, MO, USA) and Ionomycin (1 µM, #I0634 Sigma-Aldrich Chemical Co.) for 6 h, then we centrifuged the cultures to take separately the supernatant and the pellet, both were stored at − 80 °C.

### mRNA extraction and RT-qPCR

PBMCs were homogenized and total RNA was extracted using PureLink RNA Mini kit (#12183018A, Invitrogen) according to manufacturer’s recommendations. RNA purity and quantity were determined with Nanodrop 1000 spectrophotometer (Thermo Fisher). 500 ng of purified RNA was reverse transcribed using the High Capacity cDNA Reverse Transcription Kit (Applied Bio systems, Cat. #436814). The cDNA synthesis was performed at 25 °C for 10 min, at 37 °C for 120 min and a final step at 85 °C for 5 min in a final volume of 20 μl as instructed by manufacturer. Then, cDNA was analyzed by quantitative RT-PCR using PrimeTime qPCR Assays (Integrated DNA Technologies, USA). Human assays: IKZF3 (Hs.PT.58.4515293), and ACTB (Hs.PT.39a.22214857). Mouse assays: IKZF3 (Hs. PT.58.2960172) and 18S (Hs.PT.39a.22214856.g). Quantitative PCR was performed in 12 μl of final volume on 96-well plates using the Premix Ex Taq (Takara Biotechnology, #RR037A). Reactions included Segment 1: 1 cycle of 30 s at 95 °C and Segment 2: 40 cycles of 5 s at 95 °C and 20 s at 60 °C. All RT-PCR assays were run in duplicate. To provide negative control and exclude contamination by genomic DNA, the PrimeScript RT enzyme was omitted in the cDNA synthesis step and samples were subjected to the PCR reaction in the same manner with each probe. ACTB (ActinB) gene expression was used as housekeeping gene. To analyze the relative changes in gene expression, the 2( ΔΔ C(T)) method was used.

### Immunoblotting

Animals were killed by cervical dislocation. The regions of interest were dissected out, frozen using CO2 pellets and stored at −80 °C until use. Briefly, the tissue was lysed by sonication in 250 μl of lysis buffer (PBS, 10 ml l−1 Nonidet P-40, 1 mM PMSF, 10 mg l−1 aprotinin, 1 mg l−1 leupeptin and 2 mg l−1 sodium orthovanadate). After lysis, samples were centrifuged at 12,000 r.p.m. for 20 min. Supernatant proteins (15 µg) from total brain region extracts were loaded in SDS–PAGE and transferred to nitrocellulose membranes (GE Healthcare, UK). Membranes were blocked in TBS-T (150 mM NaCl, 20 mM Tris–HCl, pH 7.5, 0.5 ml Tween 20) with 50 g l − 1 non-fat dry milk and 50 g l − 1 BSA. Immunoblots were probed with the following antibody: anti-Ikzf3 (1:1000, Proteintech, #19055-1-PBS). All blots were incubated with the primary antibody overnight at 4 °C by shaking in PBS with 0.2 g l − 1 sodium azide. After three washes in TBS-T, blots were incubated for 1 h at room temperature with the corresponding horseradish peroxidase-conjugated antibody (1:2000; Promega, USA) and washed again with TBS-T. Immunoreactive bands were visualized using the Western Blotting Luminol Reagent (Santa Cruz Biotechnology, #sc-2048) and quantified by a computer-assisted densitometer (Gel-Pro Analyzer, version 4, Media Cybernetics). For loading control, a mouse monoclonal antibody for actin was used (1:20,000; MP Biochemicals, #0869100-CF).

### Animals

In the present study, all the WT mice corresponded to the strain C57BL/6 (MGI catalog #5657800, RRID: MGI:5657800), and mice from this line were backcrossed to generate the transgenic line used. Heterozygous Thy1-EGFP transgenic mice were used for dendritic spine analyses. In this line, EGFP is expressed under the neuronal Thy1 promoter in a subset of projection neurons, resulting in sparse fluorescent labeling that enables high-resolution imaging of dendritic arbors and individual spines. The Aiolos knockout (Aio^-/-^) mouse line used in this investigation has been previously described elsewhere [24]. The animals were housed with access to food and water ad libitum in a colony room kept at 19–22 °C and 40–60% humidity, under an inverted 12/12 h light/dark cycle (from 8 A.M. to 8 P.M.). All animal procedures were approved by local committees [Universitat de Barcelona, CEEA (136/19); Generalitat de Catalunya (DAAM 10786) following the European Communities Council Directive (86/609/EU).

### LPS mouse model

WT Pups at P1 and at P2 were intraperitoneally injected with 6 mg/kg of LPS or vehicle. Twenty-four hours after the second injection mice were sacrificed, brains were collected, and the hippocampus was dissected and frozen at –80 for posterior analysis. This mouse model was approved by local committees (University of Barcelona, 225/17 and Generalitat de Catalunya, 404/18), in accordance with the European Communities Council Directive (86/609/EU).

### Stereotaxic surgeries

AAVs injection: During isoflurane anesthesia (2% induction, 1.5% maintenance; 2% oxygen), we performed bilateral hippocampal injections of either AAV8-CamKII-m-IKZF3-P2A-mCherry or AAV8-CamKII-mCherry. We used the following coordinates (millimeters) from bregma (anteroposterior and lateral) and from skull (dorsoventral); anteroposterior: −2.0; Lateral +/−1.3, and dorsoventral: −1.4 (CA1) and −2 (Dentate Gyrus). The cannula was left to deliver 0.75 μl of 1:1 virus in each depth during 2 minutes and five additional minutes were left to have complete virus diffusion. After 2 h of monitoring, mice were returned to their home cage for 4 weeks before the next experiments.

Mini-osmotic pumps implantation: after isoflurane anesthesia (2% induction, 1.5% maintenance; 2% oxygen), mice were placed in a stereotaxic apparatus for mini osmotic pump (model 1004; Alzet, Palo Alto, CA, USA) implantation. We used a brain infusion kit (#0008663) to deliver into the lateral (left) ventricle (0.1 mm posterior to bregma, ± 0.8 mm lateral to the midline, and − 2.5 mm ventral to the parenchyma surface) 0.11 μl per hour of supernatant obtained from cultured peripheral blood mononuclear cells (PBMCs) at a protein concentration of 2,064 µg/µl. Cannulas were fixed on the skull with Loctite 454 (from Alzet). Minipumps, previously equilibrated overnight at 37 °C in PBS, were implanted subcutaneously in the back of the animal. After recovery, the mice were isolated to prevent detachment of the cannulas.

### Behavioral tests

Behavioral phenotyping was performed following widely and previously detailed features of appropriate mouse models of schizophrenia addressing the three dimensions of the symptomatology [25].

Locomotion and evaluation of Ketamine sensitivity (Ketamine, 30 mg/kg; TOCRIS 2813), were both evaluated in the open field arena (a white square arena measuring 40 × 40 × 40 cm in length, width, and height respectively). For locomotion, mice were left in the middle of the arena, and their behavior was recorded for 10 minutes. For Ketamine sensitivity, mice were left to habituate for 4 minutes. After this period, mice received an intraperitoneal injection of vehicle (0.9% NaCl), or Ketamine and their behavior was recorded for 10 minutes. Dim light intensity was 60 lux throughout the arena.

Next, sociability was assessed by using the three-chamber social interaction test as previously described [9]. Finally, the cognitive domain was evaluated with the novel object recognition test (NORT) and the novel object location test (NOLT) as previously reported [8], [26]. In all behavioral experiments, animals were tracked and recorded with Smart junior 3.0 software (Panlab).

### Quantitative proteomics

Protein quantitation was performed with the Brad ford assay kit (Bio-Rad, Hercules, CA, USA). Protein extracts (20 µg) were diluted in Laemmli sample buffer and loaded into a 1,5 mm thick polyacrylamide gel with a 4% stacking gel cast over a 12.5% resolving gel. The run was stopped as soon as the front entered 3 mm into the resolving gel so that the whole proteome became concen trated in the stacking/resolving gel interface. Bands were stained with Coomassie Brilliant Blue, excised from the gel and protein enzymatic cleavage was carried out with trypsin (Promega; 1:20, w/w) at 37 °C for 16 h as previously described [27]. Purification and concentration of peptides were performed using C18 Zip Tip Solid Phase Extraction (Millipore). For LC–MS/MS, dried peptide samples were reconstituted with 2% ACN-0.1% FA (Ace tonitrile-Formic acid) and quantified by NanoDropTM spectrophometer (ThermoFisher) before LC–MS/MS analysis using an EASY-1000 nanoLC system coupled to an Exploris 480 mass spectrometer (Thermo Fisher Sci entific). Peptides were resolved using a C18 Aurora col umn (75 µm × 25 cm, 1.6 µm particles; IonOpticks) at a flow rate of 300 nl/min using a 60-min gradient (50 °C): 2% to 5% B in 1 min, 5% to 20% B in 48 min, 20% to 32% B in 12 min, and 32% to 95% B in 1 min (A = FA, 0.1%; B = 100% ACN:0.1% FA). The spray voltage was set at 1.6 kV, and the capillary temperature at 275 °C. Sample data was acquired in a data-independent mode (DIA) with a full MS scan (scan range: 400 to 900 m/z; resolu tion: 60,000; maximum injection time: 50 ms; normalized AGC target: 100%) and 25 periodical MS/MS segments, applying 20 Th isolation windows (0.5 Th overlap; reso lution: 15,000; maximum injection time: 22 ms; normal ized AGC target: 3000%). Peptides were fragmented using a normalized HCD collision energy of 30%. Data were acquired in profile and centroid mode for full MS scan and MS/MS, respectively. For data analysis of quantitative proteomics DIA data files were analyzed using Spectronaut (v 17.3, Biognosys) by directDIA analy sis (dDIA). Sample raw data was first processed by the Pulsar Spectronaut search engine to generate a spec tral library. Pulsar then searched MSMS spectra against Uniprot-Swissprot isoforms Homo Sapiens database. MS1/MS2 calibration and main search tolerance were set to dynamic. The maximum precursor ion charge was set to 4, and fragment selection to intensity-based. Car bamidomethyl (C) was selected as a fixed modification, and Oxidation (M), Acetyl (Protein N-term), Deamida tion (N), and Gln-> pyro-Glu as variable modifications (3 maximum modifications per peptide). The enzyme was set to trypsin in a specific mode (two missed cleav ages maximum). The target-decoy-based false discovery rate (FDR) filter for PSMs, peptide, and protein groups was set to 1%. Once the Pulsar search was performed, the Spectronaut DIA search engine algorithm initiated the DIA search using the default settings (Proteotypicity filter = Protein group Specific) and filtering the precur sors and protein groups by a 1% Q-value. The Perseus software (version 1.6.15.0) was used for statistical analysis and data visualization [28]. Identifications from the reverse database, common contaminants and proteins only identified through a modification peptide were removed. Label-free intensities were then logarithmized (base 2) and the samples were then grouped according to the experimental design. A filter of 70% valid values at least in one group was applied to the resulting matrix followed by a logarithmic transformation (Log2). Then, data were imputated based on the normal distribution and normalized using the “Width adjustment” method. Sta tistical analysis was performed using “Two-sample tests” for the T-test of two selected experimental groups, based on permutation-based FDR statistics (250 permutations; FDR 1⁄4 0.07; S0 1⁄4 0.1). The resulting matrixes were exported containing the p-value (-Log) and fold-change (Log2) columns that, after its transformation into linear scale, were filtered by p < 0.05 and 30% of significance and fold-change respectively.

### Hippocampal primary try-cultures and immunofluorescence

Tri-cultures were prepared from the hippocampi of P0 WT mice, following a protocol from a previously published study with minor adaptations [29]. For immunofluorescence, cells were fixed for 10 min with 40 g l−1 paraformaldehyde in PB 0.2 M for immunostaining. Fixed cells were permeabilized in 1 ml l−1 Triton X-100 for 10 min and then blocking was performed with 10 g l−1 BSA in PBS for 1 h. Cells were incubated with mouse monoclonal antibodies anti-Iba1 (1:500, Abcam, #Ab5076), anti-GFAP (1:500, Dako, #Z0334) and anti-iC3b (1:200, Gene Tech, # GTX40522) at 4 °C overnight. After three washes with PBS, cells were incubated with the corresponding fluorescent secondary antibodies Alexa Fluor 488, 555 or 647 (1:400; from Jackson Immuno Research, West Grove, USA). Nuclei were stained with DAPI Fluoromount (SouthernBiotech, USA). Cells were analyzed using a two-photon confocal microscope (Leica SP5).

### Tissue fixation and immunofluorescence

Mice were euthanized by cervical dislocation. Left hemispheres were removed and fixed for 72 h in 4% paraformaldehyde (PFA) in PBS. Thy1-EGFP mice were anaesthetized (pentobarbital, 60 mg kg−1 i.p.) and intracardially perfused with a 40 g l−1 paraformaldehyde solution in 0.1 M sodium phosphate, pH 7.2. Then brains were removed and fixed for 72 h in 4% paraformaldehyde (PFA) in PBS. 40 µm or 70 µm (for Thy1-EGFP mice) coronal sections were obtained using a Leica vibratome (Leica VT1000S). Next, free-floating sections were washed three times in PBS, treated with NH4Cl for 30 min, and washed again three times with PBS. Floating sections were permeabilized and blocked with PBS containing 3% Triton X-100, 0.02% Azide, 2% BSA, and 3% NGS (Ab buffer) for 1 h at RT. After three washes in PBS, brain slices were incubated overnight at 4 °C with the following antibodies anti-Ikzf3 (1:200, Genetech, #gtx64869), anti-Iba1 (1:500, Abcam, #Ab5076), anti-GFAP (1:500, Dako, #Z0334), anti-NeuN (1:1000, Chemicon, Ref: MAB377), anti-GFP (1:500, Synaptic Systems, 132006), anti-PSD-95 (1:500, Cell Signaling, #3450S), anti-iC3b (1:200, Gene Tech, # GTX40522). Sections were washed three times and incubated for 2 h at RT with fluorescent secondary antibody Alexa Fluor 488, 555 or 647 (1:400; from Jackson Immuno Research, West Grove, USA). Nuclei were stained with DAPI Fluoromount (SouthernBiotech, USA). Sections were analyzed using a two-photon confocal microscope (Leica SP5).

### Imaging analysis

All immunostained materials, cells and tissues were imaged using a Leica Confocal SP5 (63 × numerical aperture lens, 5 × digital zoom, 1-Airy unit pinhole). Confocal z-stacks were taken at 1,024 × 1,024 pixel resolution, every 1 μm for brain sections and every 0.2 μm for cultured cells and spine analysis. Subsequent analyses were performed with NIH ImageJ freeware (Wayne Rasband, NIH). To analyze labelled iC3b-positive puncta in vitro we selected dendritic segments with clear visualization. No more than one per neuron and at least 40 per group, the total number of iC3b-positive puncta was obtained using the cell counter tool in the ImageJ software. For spines counting, we selected dendritic segments of secondary dendrites 45 μm away from the cell body. The total number of spines was obtained using the cell counter tool in the ImageJ software. At least 8 dendrites per animal from 5 mice per group were counted. For the experiments evaluating PSD95, iC3b and Iba1 in vivo we directly counted positive clusters per field using a homemade algorithm in ImageJ software. At least 8 fields per animal from 5 mice per group were counted.

### Statistics

All data are expressed as mean ± SEM. Normal distribution was tested with the D’Agostino and Pearson omnibus test. If no difference in normal distribution was detected, statistical analysis was performed using two-tailed Student’s t test (95% confidence) or ANOVA and Tukey’s or Dunnett’s post hoc tests. If distribution was not normal, nonparametric two-tailed Mann–Whitney test was used. In case of unequal variances in the t-test the Welch’s correction was applied. Values of p < 0.05 were considered statistically significant. In Figure 1, due to the small sample size, normality could not be reliably assessed, and differences among groups were analyzed using one-way ANOVA and two-tailed Student’s t test. All experiments in this study were blinded and randomized. All mice bred for the experiments were used for preplanned experiments and randomized to experimental groups. Visibly sick animals were excluded before data collection and analysis. Data was collected, processed, and analyzed randomly. The experimental design and handling of mice were identical across experiments. Littermates were used as controls with multiple litters (3–5) examined per experiment.

**Figure 1.**
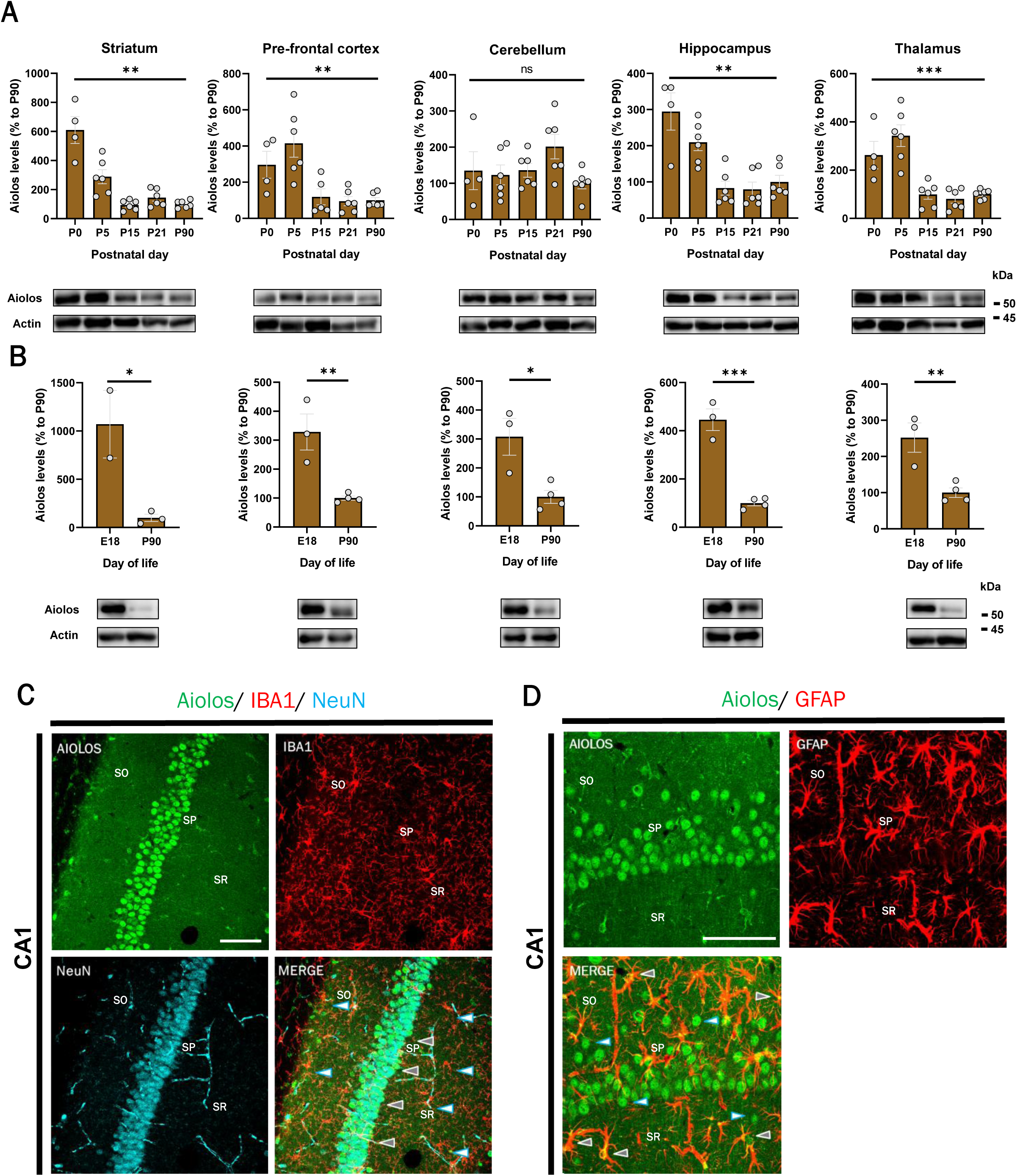
Developmental expression and cellular localization of Aiolos in WT mouse brains. (A) Aiolos protein levels were assessed by Western blot in the striatum, prefrontal cortex, cerebellum, hippocampus, and thalamus at postnatal days 0 (P0), 5 (P5), 15 (P15), 21 (P21), and 90 (P90). (B) Comparison of Aiolos protein levels between embryonic day 18 (E18) and P90 in the same brain regions. Representative Western blots are shown below the corresponding graphs. Aiolos protein levels were normalized to actin and expressed relative to P90. Each dot represents an individual animal, with males and females equally represented. Data are presented as mean ± SEM. Statistical differences were assessed using one-way ANOVA (A) or two-tailed Student’s t-test (B). ns, not significant; *\** p < 0.05, ** p < 0.01, ***p < 0.001. (C–D) Representative immunofluorescence images showing Aiolos expression in the CA1 region of the hippocampus in adult WT mice (>P90). (C) Immunostaining for Aiolos (green), Iba1 (red), and NeuN (cyan). Gray arrowheads indicate Aiolos expression in neurons, while white arrowheads indicate macrophages. (D) Immunostaining for Aiolos (green) and GFAP (red). Gray arrowheads indicate Aiolos immunoreactivity in astrocytes, while white arrowheads indicate Aiolos expression in other cell populations. SO, stratum oriens; SP, stratum pyramidale; SR, stratum radiatum. Scale bars: 100 µm.

## RESULTS

### 1. Aiolos displays a differential protein expression dynamic through development in different mouse brain regions and stabilizes in adulthood, with predominant expression in principal neurons in the hippocampus

As it has been previously shown that Ikaros and Helios can be differentially expressed during neurodevelopment in different brain regions [6], [7], [30], Our first aim was to characterize Aiolos expression at this stage. We explored Aiolos protein levels throughout development across different brain regions in WT mice. Aiolos expression was assessed at E18, P0, P5, P15, P21, and P90 in the striatum, prefrontal cortex, cerebellum, hippocampus, and thalamus. We found that Aiolos protein levels changed significantly between P0 and P90 in all regions analyzed, with the exception of the cerebellum (Figure 1A). Furthermore, in all brain regions examined, Aiolos levels were lower at P90 than at E18 (Figure 1B), where expression levels appeared to stabilize. Subsequently, we aimed to explore in which type of neural cell Aiolos is expressed. Using immunofluorescence in adult WT mouse brains, we observed prominent nuclear expression of Aiolos in hippocampal principal neurons, together with an apparent cytoplasmic Aiolos immunoreactivity in a subset of astrocytes (Figure 1C-D).

### 2. Knocking down Aiolos in mice generates minor changes in glia cells and a specific molecular profile in the hippocampus

Previous studies using Ikaros and Helios knockout (KO) models have provided insights into their functions in the adult mouse brain [8], [9]. Therefore, after characterizing the protein expression pattern of Aiolos, we next investigated whether its deletion induced cellular alterations in the adult mouse brain. Using immunofluorescence analysis, we assessed morphological parameters of astrocytes and microglia in the hippocampus. GFAP-positive cells from Aiolos KO mice exhibited an increase in cell area and minimum feret diameter compared with WT mice (Figure 2B). On the other hand, IBA1-positive cells did not exhibit alterations in any of the parameters analyzed (Figure 2C). Furthermore, as Aiolos is expressed in neurons, we also assessed whether its absence affected neuronal abundance. Consistent with the lack of major cellular alterations, Aiolos KO mice did not show differences in NeuN-positive cells density compared with WT mice (Figure 2D). To further characterize the molecular consequences of Aiolos deficiency, we sought to determine whether the absence of Aiolos was associated with alterations in hippocampal protein expression. To this end, we performed a mass spectrometry-based proteomic analysis using hippocampal tissue from Aiolos KO and matched wild type (WT) mice. A total of 95 differentially expressed proteins (DEPs) were identified, including 45 upregulated and 50 downregulated proteins in Aiolos KO mice compared with CNT mice (Figure 2E). Among the downregulated proteins C1qa and C1qb were identified. Whereas H4c1, Flrt3 and Wdfy1 were found upregulated. To gain insight into the biological processes associated with the identified DEPs, we subsequently examined the reported functions of the differentially expressed proteins. Through literature-based analysis, we observed that multiple altered proteins, such as C1qa, C1qb and Wdfy1 among others, were directly or indirectly related to schizophrenia [9], [31], [32], suggesting that Aiolos could by implicated in schizophrenia pathology-related pathways.

**Figure 2.**
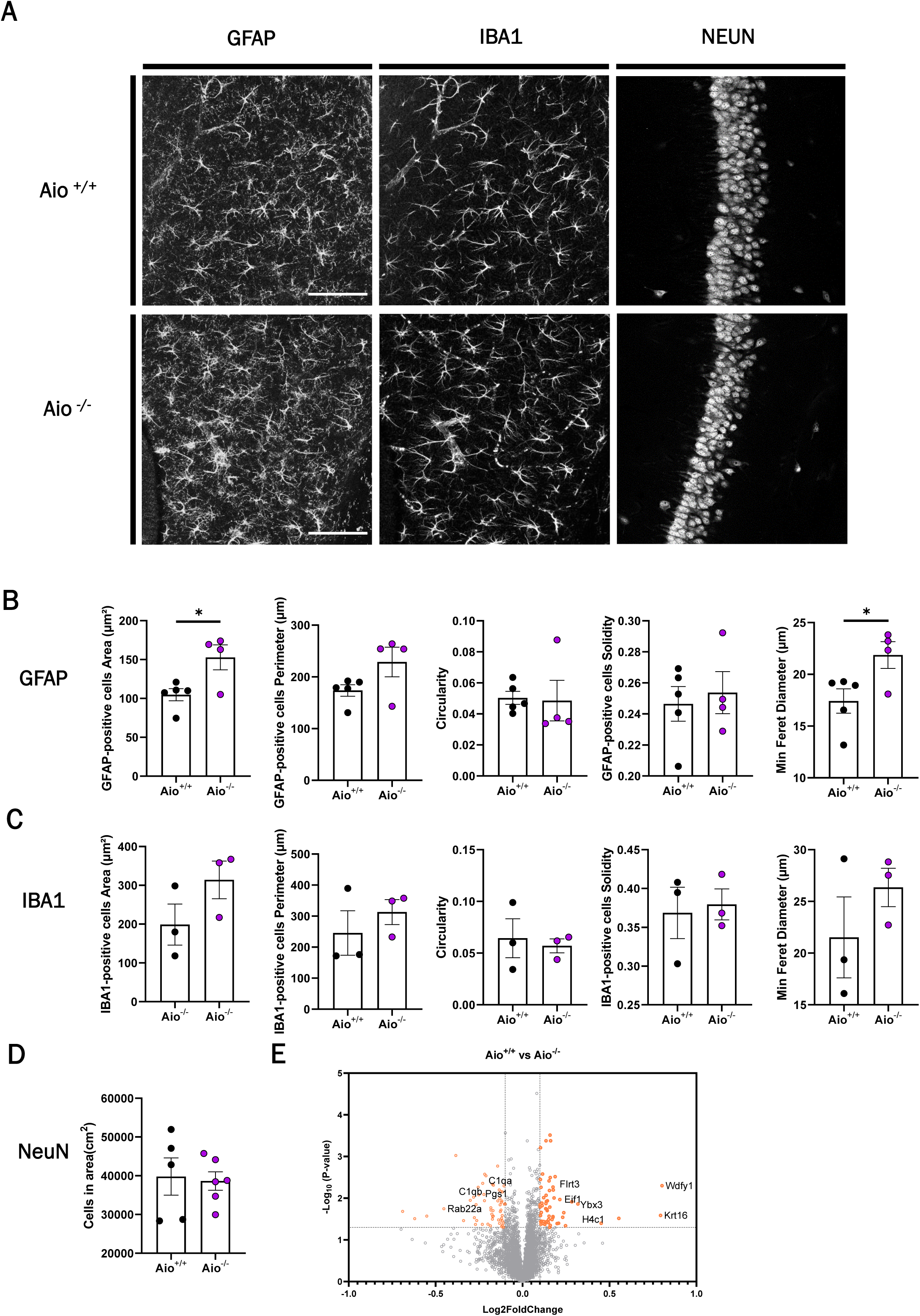
Cellular and proteomic alterations in the hippocampus of Aiolos KO mice. (A) Representative immunofluorescence images showing GFAP-positive astrocytes, IBA1-positive microglia, and NeuN-positive neurons in the CA1 region of the hippocampus from adult (>P90) WT (Aio⁺/⁺) and Aiolos KO (Aio⁻/⁻) mice. Scale bars: 100µm. (B–C) Quantification of morphological parameters, including cell area, perimeter, circularity, solidity, and minimum Feret diameter, in GFAP-positive astrocytes (B) and IBA1-positive microglia (C). (D) Quantification of NeuN-positive cell density in the hippocampus. (E) Volcano plot showing differentially expressed proteins identified by mass spectrometry-based proteomic analysis of hippocampal tissue from Aio⁺/⁺ and Aio⁻/⁻ mice. The x-axis represents log₂ fold change and the y-axis represents −log₁₀(*p*-value). Green dots indicate significantly differentially expressed proteins, while dark dots represent non-significant changes. Selected proteins are labeled in red. Proteomic statistical analysis was performed using a two-sample t-test with permutation-based FDR correction (250 permutations, FDR = 0.07, S0 = 0.1), followed by filtering at *p* < 0.05 and a 30% fold-change threshold. Data in panels B–D are presented as mean ± SEM, with each dot representing an individual animal. Statistical differences in panels B–D were assessed using two-tailed Student’s t-tests. * < 0.05.

### 3. Aiolos expression is increased in PBMCs and the Hippocampus of schizophrenia patients and in a mouse model relevant to schizophrenia while it is not altered in patients with major depressive disroder

Given that several of the DEPs identified in our proteomic analysis have previously been associated with schizophrenia, and considering that Ikaros and Helios, have also been implicated in schizophrenia-related phenotypes [9], [10], we next aimed to investigate the potential involvement of Aiolos in schizophrenia. First, as Ikaros and Helios have been shown to be dysregulated in PBMCs from schizophrenia patients [10], we evaluated Aiolos levels in the same cell population. Notably, Aiolos was significantly upregulated in PBMCs from schizophrenia patients compared to controls (Figure 3A). We then assessed whether Aiolos dysregulation was also present in the brain and found increased Aiolos levels in the hippocampus (post-mortem) of schizophrenia patients relative to matched controls (Figure 3B). To assess the specificity of these findings, we next evaluated Aiolos levels in patients with Major Depressive Disorder (MDD). In contrast to schizophrenia, Aiolos levels were unaltered in both PBMCs and hippocampal post-mortem samples from MDD patients compared with controls (Figure 3C-D), suggesting that Aiolos dysregulation may be specific to schizophrenia. Finally, to further investigate the involvement of Aiolos in the context of schizophrenia, we evaluated Aiolos levels in the hippocampus of an animal model relevant to schizophrenia, the neonatal Lipopolysaccharides (LPS) model [33]. We observed that Aiolos was upregulated in the hippocampus of the animals treated with LPS compared with the vehicle-treated ones (Figure 3E-F), strengthening the idea of the involvement of Aiolos in Schizophrenia.

**Figure 3.**
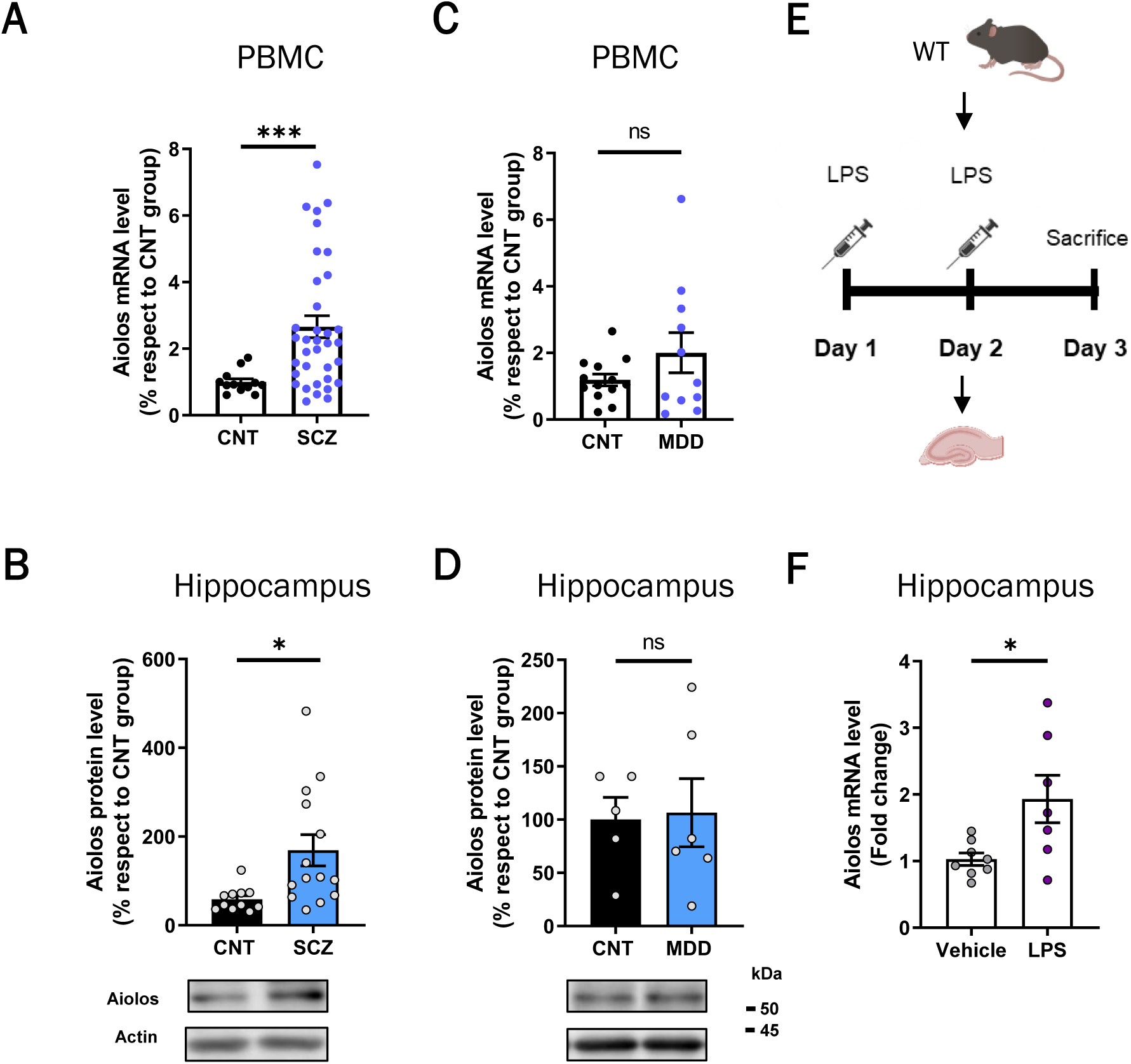
Aiolos upregulation in schizophrenia patients and a neonatal LPS mouse model. (A–B) Aiolos mRNA levels measured by RT-qPCR in peripheral blood mononuclear cells (PBMCs) (A) and protein levels assessed by Western blot in hippocampal tissue (B) from schizophrenia (SCZ) patients and healthy controls (CNT). (C–D) Aiolos mRNA levels measured by RT-qPCR in PBMCs (C) and protein levels assessed by Western blot in hippocampal tissue (D) from major depressive disorder (MDD) patients and healthy controls (CNT). Aiolos expression levels are presented relative to the corresponding control groups. (E) Schematic representation of the neonatal lipopolysaccharide (LPS) administration protocol in WT mice, consisting of intraperitoneal injections on postnatal days 1 and 2, followed by sacrifice and hippocampal tissue collection on day 3. (F) Aiolos mRNA levels measured by RT-qPCR in hippocampal tissue from LPS-treated and vehicle-treated mice. Data are presented as mean ± SEM, with each dot representing an individual patient or animal. Statistical differences were assessed using the Mann–Whitney U test (A, B, D, F) or unpaired Student’s t-test (C). ns, not significant; *\** < 0.05, ** < 0.001.

### 4. Induced overexpression of Aiolos in the hippocampus of WT mice generates positive and cognitive schizophrenia-like symptoms

To investigate the consequences of hippocampal Aiolos upregulation in schizophrenia, we performed a model of Aiolos overexpression in the hippocampus of WT mice (Aiolos OV). Adult WT mice received hippocampal injections of an Adeno associated virus (AAV) vector expressing Aiolos under the control of the CamKIIα promoter. We injected AAV8-CamKII-m-IKZF3-P2A-mCherry (AAV-Aiolos) to overexpress Aiolos or AAV8-CamKII-mCherry (AAV-mCherry) as a control (Figure 4A). Immunofluorescence analysis confirmed successful transduction and viral expression in the dentate gyrus (Figure 4B). We next hypothesized that Aiolos overexpression would induce behavioral phenotypes relevant to schizophrenia. Consistently, mice receiving AAV-Aiolos exhibited increased locomotor activity and a higher parallel index in the open field test compared with control animals (Figure 4C), indicative of a hyperactive phenotype (mimicking agitation). Furthermore, following ketamine administration, AAV-Aiolos mice displayed a reduction in the parallel index relative to controls (Figure 4D), suggesting an increased sensitivity to the behavioral effects of psychotomimetic drugs. We next assessed cognitive performance using the Novel Object Location Test (NOLT). Control animals preferentially explored the object in the novel location, whereas AAV-Aiolos mice failed to discriminate between the familiar and the novel location (Figure 4E), suggesting a cognitive deficit following Aiolos overexpression.

**Figure 4.**
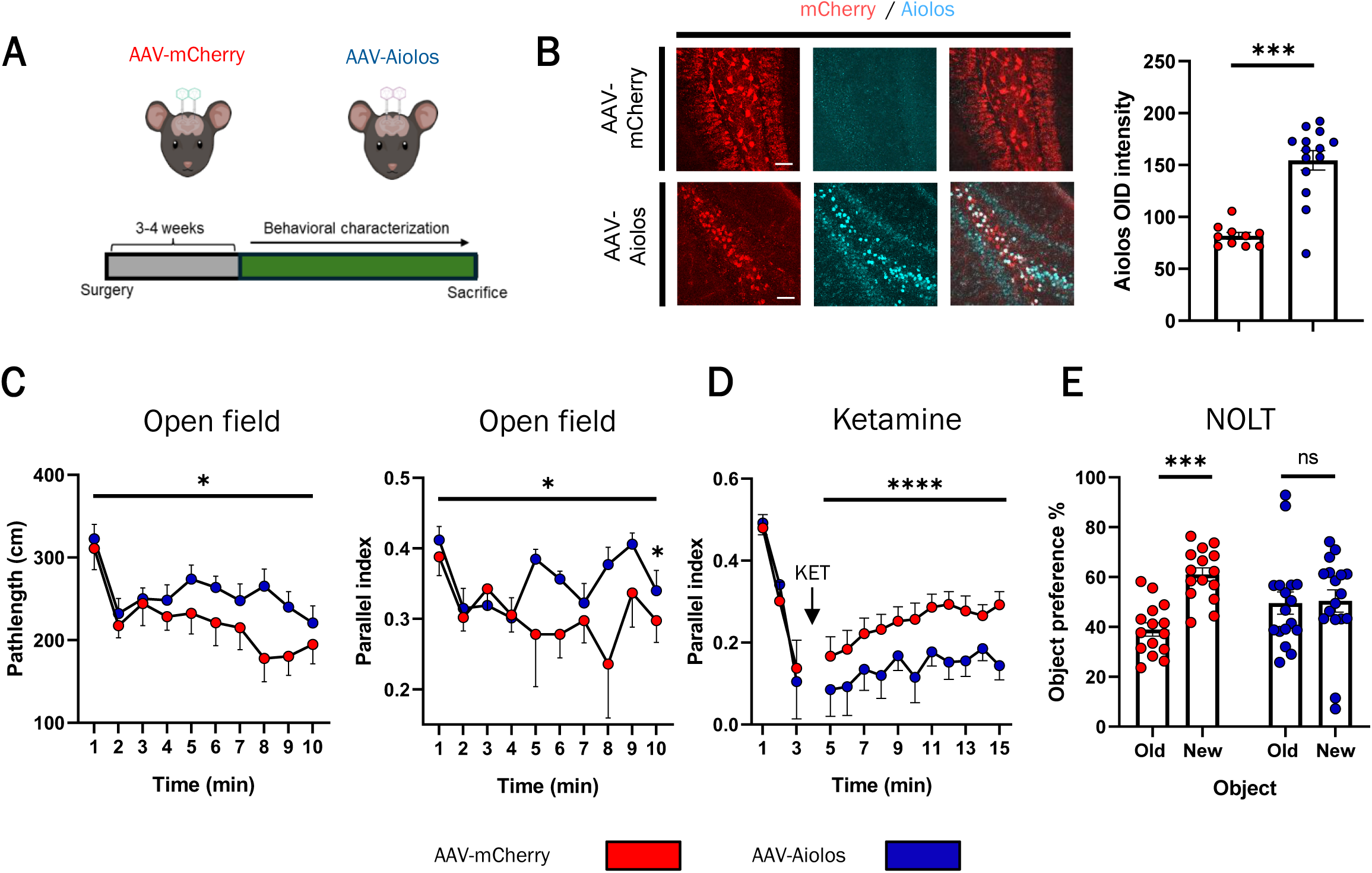
Hippocampal Aiolos overexpression induces behavioral alterations in WT mice. (A) Schematic representation of the experimental design. Adult WT mice received hippocampal injections of adeno-associated viral (AAV) vectors expressing Aiolos/mCherry (AAV-Aiolos) or mCherry (AAV-mCHERRY). Behavioral characterization was performed 3–4 weeks after surgery. (B) Representative immunofluorescence images and quantification of Aiolos fluorescence intensity in the dentate gyrus of AAV-Aiolos mice compared with AAV-mCHERRY controls. Scale bars: 50 µm. (C) Locomotor activity, assessed by path length, and parallel index measured during the open field test. (D) Parallel index measured following ketamine administration. (E) Percentage of time spent exploring familiar (old) and novel (new) object location during the novel object location test (NOLT). Data are presented as mean ± SEM, with each dot representing an individual animal. In panel E, familiar and novel object exploration values were obtained from the same animals. Statistical differences were assessed using unpaired Student’s t-tests (C–D) or two-way ANOVA followed by Sidak’s multiple comparisons test (E). ns, not significant; *p < 0.05, ***p < 0.001, ****p < 0.0001.

### 5. PBMCs secretome of schizophrenia patients with upregulated levels of Aiolos generates negative schizophrenia-like symptoms in WT mice

After investigating the consequences of hippocampal upregulation of Aiolos. We aimed to assess the possible effects of the immune dysregulation of Aiolos. To this end, we used a translational model. We obtained and cultured PBMCs from schizophrenia patients and controls and obtained their supernatants containing the secretome of the cells (Figure 5A). We specifically selected patients with an up-regulation of Aiolos (> 40%) in PBMCs compared to the control subjects (Figure 5B), both groups were matched in terms of number (n = 5/group), age and sex (Supplementary Table 5). Next, we generated two pools with the secretomes, one corresponding to the group of schizophrenia patients and the other one to the group of control subjects and then, we treated adult WT mice with the secretomes. We injected directly and permanently the secretomes into the lateral ventricles of the animals using mini-osmotic pumps at a delivery ratio of 2,5 µl/day for 28 days (Figure 5C). Four days after the surgery, we performed behavioral testing (Figure 5D). When evaluated in the open field test, no differences in locomotor activity were observed between the two groups (Figure 5E). Similarly, when the cognitive performance was evaluated in the Novel Object Recognition Test (NORT), both groups successfully recognized the novel object (Figure 5F). In contrast, mice receiving secretome derived from schizophrenia patients displayed altered social behavior in the three-chamber sociability test (TCT), whereas mice receiving secretome from control subjects did not (Figure 5G), indicating a specific social behavior alteration induced by the schizophrenia patients derived secretome obtained from their PBMCs.

**Figure 5.**
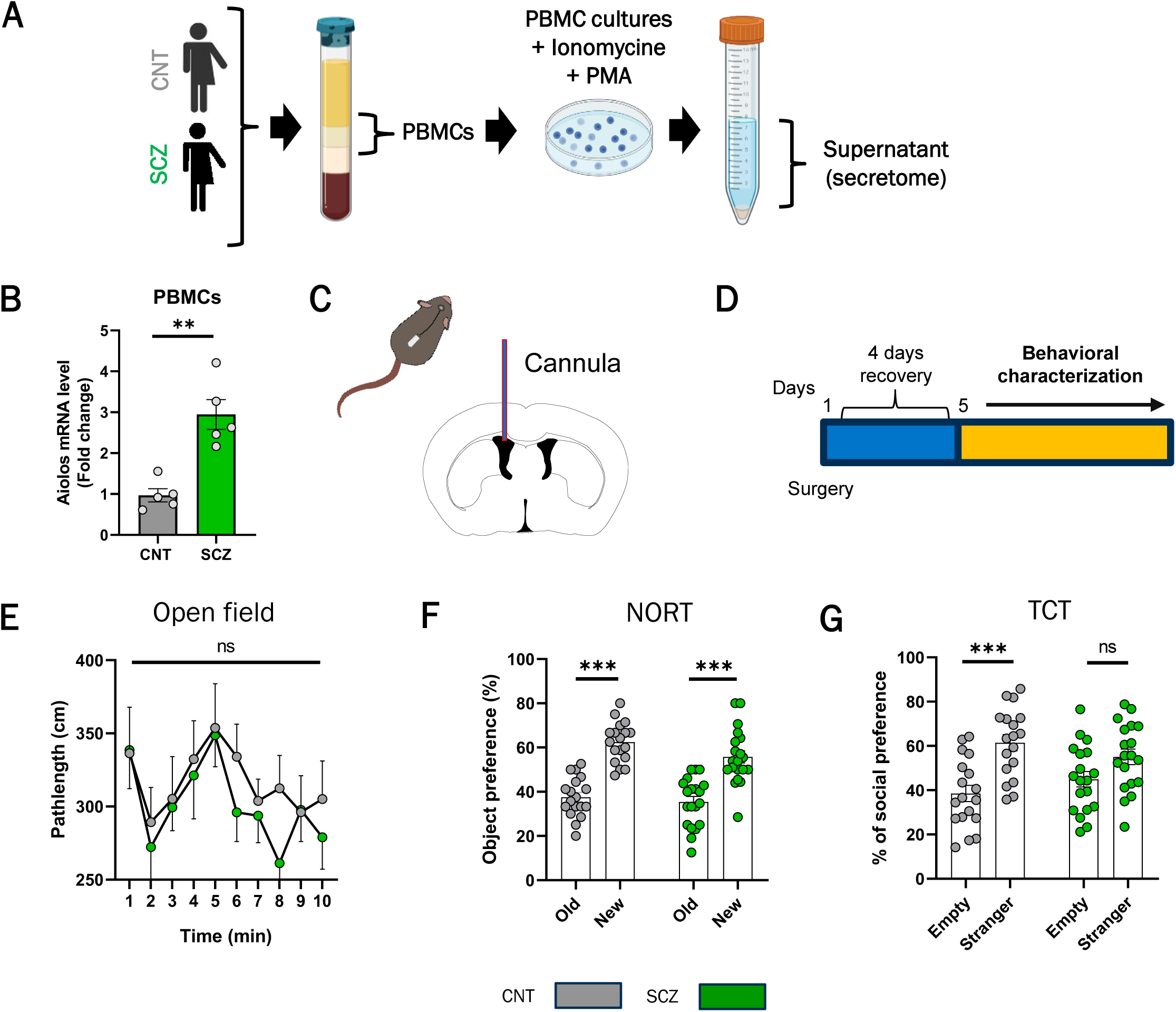
Behavioral characterization of WT mice receiving PBMC-derived secretomes from schizophrenia patients with elevated Aiolos expression. (A) Schematic representation of secretome preparation. PBMCs from control subjects (CNT) and schizophrenia patients (SCZ) were stimulated with PMA and ionomycin for 6 h before supernatant collection. Equal volumes from five donors per group were pooled to generate control and SCZ secretomes. (B) Relative Aiolos mRNA levels measured by RT-PCR in PBMCs from control subjects and selected schizophrenia patients. (C) Schematic representation of intracerebroventricular secretome delivery into adult WT mice using mini-osmotic pumps at a rate of 2.5 µL/day for 28 days. (D) Experimental timeline showing surgery, four days of recovery, and subsequent behavioral characterization. (E) Locomotor activity, assessed by path length during the open field test. (F) Percentage of time spent exploring familiar (old) and novel (new) objects during the novel object recognition test (NORT). (G) Social preference during the three-chamber sociability test (TCT). Data are presented as mean ± SEM. Each dot represents an individual donor in B or an individual animal in F–G. Familiar and novel object exploration values (F), and empty and stranger exploration values (G), were obtained from the same animals. Statistical differences were assessed using Welch’s t-test (B), two-way ANOVA (E), or two-way ANOVA followed by Bonferroni’s multiple comparisons test (F–G). ns, not significant; **p < 0.01, ***p < 0.001.

### 6. Different proteomic profiles pinpoint complement dysregulation as a candidate mechanism of Aiolos action in the hippocampus

Considering that the mouse model mimicking the alteration of Aiolos in the hippocampus showed the strongest phenotype, we aimed to further investigate the implication of Aiolos in this context. First, we sought to determine whether the overexpression of Aiolos was associated with alterations in hippocampal protein expression. By performing a mass spectrometry-based proteomic analysis using hippocampal tissue from mice receiving AAV-Aiolos and AAV-mCherry, we identified a total of 244 DEPs, including 150 upregulated and 94 downregulated (Figure 6A). Among the downregulated proteins were Rnf112 and Eef2k whereas C1qa, C1qb, C1qc and C4b were among the proteins found to be upregulated. Posterior literature-based analysis of the DEPs revealed that several of the altered proteins have previously been linked, either directly or indirectly, to schizophrenia or schizophrenia-associated mechanisms [32], [34], [35], [36]. These findings led us to hypothesize that Aiolos could be mechanistically involved in pathways relevant to schizophrenia. Subsequently, Gene Ontology annotations (GO-Biological Process and GO-Cellular Component) and Kyoto Encyclopedia of Genes and Genomes (KEGG) were consulted to explore the biological functions associated with the DEPs. This inspection indicated that several altered proteins were associated with GO categories including, “phagocytosis” and “glial cell projection” and a KEGG category including “complement and coagulation cascades” (Figure 6C). To further investigate the relevance of Aiolos modulation in these pathways, we compared the proteomic profiles obtained from the Aiolos KO and Aiolos OV models (Figure 6B). Remarkably, we identified opposing expression patterns for members of the complement system, with C1qa and C1qb being downregulated in the Aiolos KO model and upregulated in the Aiolos OV model, suggesting that Aiolos could be essential in relation with mechanisms associated with complement proteins regulation.

**Figure 6.**
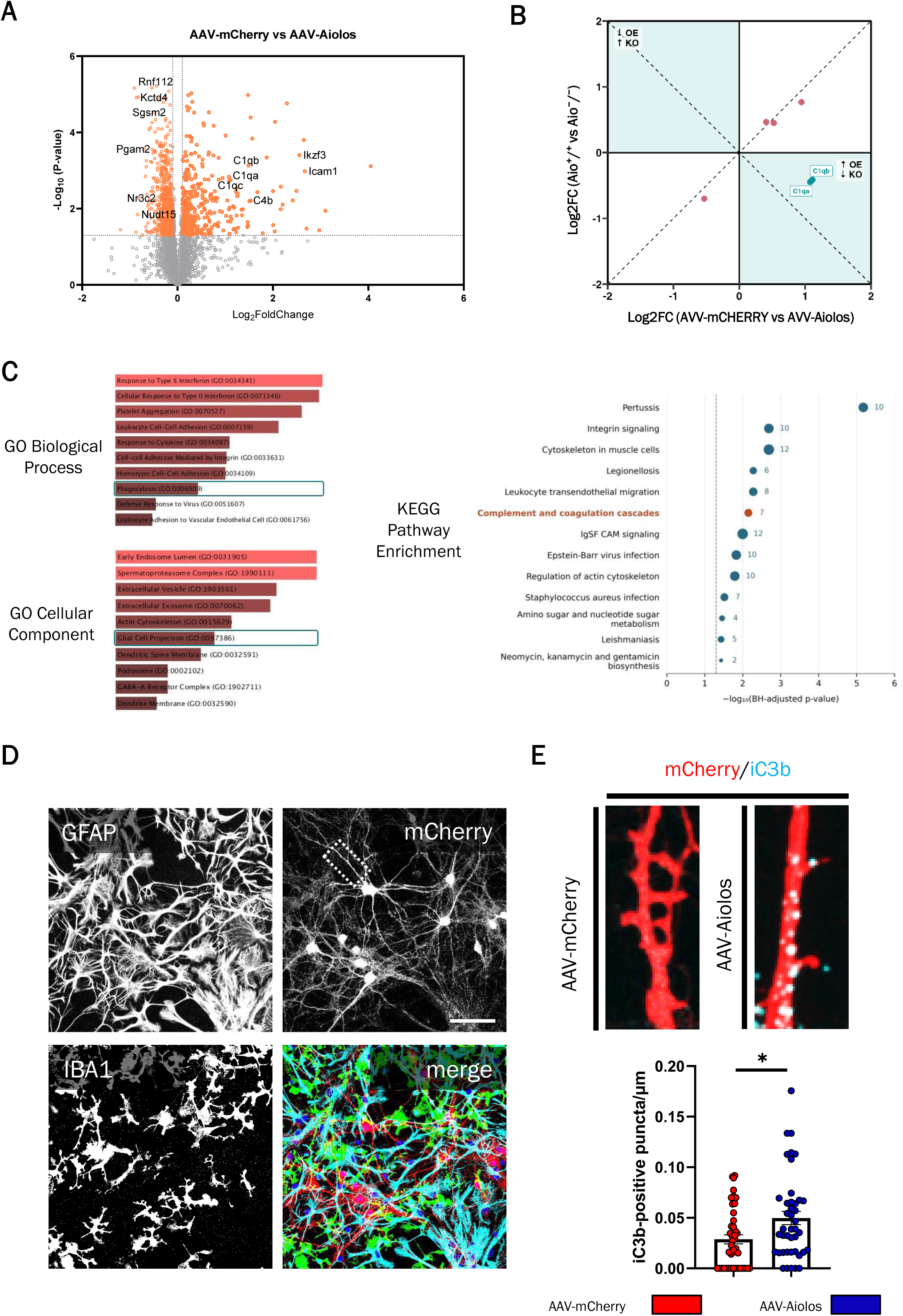
Aiolos overexpression alters hippocampal complement protein abundance and increases dendritic iC3b puncta density in vitro, with GO and KEGG enrichment of proteomic changes. (A) Volcano plot of mass spectrometry–based proteomic analysis of hippocampal tissue from mice receiving AAV-Aiolos or AAV-mCHERRY. The x-axis shows log₂ fold change and the y-axis shows −log₁₀(*p* value). Orange dots indicate the 244 differentially expressed proteins, and dark dots indicate nonsignificant changes. Selected proteins are labeled. Proteins were analyzed using a two-sample *t*-test with permutation-based false discovery rate correction (250 permutations, FDR = 0.07, S₀ = 0.1), followed by filtering at *p* < 0.05 and a 30% fold-change threshold. (B) Comparison of protein log₂ fold changes in the Aiolos knockout (KO) and overexpression (OE) models. Green dots highlight C1qa and C1qb, which were downregulated in the KO model and upregulated in the OE model. (C) The left plots show Enrichr results for GO Biological Process (top) and GO Cellular Component (bottom). Teal outlines highlight phagocytosis (GO:0006909) and glial cell projection (GO:0097386). The right plot shows the 13 KEGG pathways enriched at a Benjamini–Hochberg false discovery rate (FDR) < 0.05, based on a hypergeometric test using all mouse genes annotated to KEGG pathways as the background. Dot position indicates −log₁₀ of the adjusted P value, and dot size and adjacent numbers indicate the number of differentially abundant proteins mapped to each pathway. The dashed line marks FDR = 0.05, and the complement and coagulation cascades pathway is highlighted in orange. (D) Representative fluorescence images of mixed cultures of neurons, astrocytes, and microglia prepared from wild-type pups at postnatal day 0. Cultures received AAV-mCHERRY or AAV-Aiolos at day in vitro (DIV) 0 and were fixed at DIV 14. Images show GFAP, Iba1, and mCherry signals and their merge. Scale bar, 10 µm. (E) Representative fluorescence images of mCherry-labeled dendrites (red) and iC3b-positive puncta (cyan), with quantification of iC3b-positive puncta per µm of dendrite. Data are presented as mean ± SEM. Each dot represents one dendrite, with data collected from three independent cultures. Groups were compared using the Mann–Whitney *U* test. \**p* < 0.05.

### 7. Aiolos overexpression leads to upregulation of the complement iC3b and microglia-associated loss of synaptic material

Based on our results, we then aimed to deepen on the evaluation of complement associated proteins following Aiolos overexpression. As a first step, we performed an *in vitro* approach; we co-cultured neurons, astrocytes and microglia coming from WT pups at P0 (Figure 6D) and we transduced these cultures either with AAV-Aiolos or AAV-mCherry at day in vitro 0. At day in vitro 14 we fixed the cultures and we evaluated the presence of Ic3b (a ligand for microglial CR3), the activated cleavage product of C3, which acts downstream in the complement cascade as an opsonin tagging targets for elimination by phagocytic cells [37], [38]. We found that the dendrites of neurons treated with AAV-Aiolos presented an increased density of Ic3b-postive puncta compared with the dendrites of neurons of the control group treated with AAV-mCherry (Figure 6E). Next, we aimed to assess possible effects of complement proteins upregulation in structural synaptic plasticity following Aiolos overexpression in an in vivo model. To this end, we injected either AAV-Aiolos or AAV-mCherry into the hippocampus of Thy1-EGFP mice, a transgenic model where EGFP expression under the Thy1 promoter allows fluorescent visualization of individual neurons and their dendritic spines (Figure 7A). We then evaluated spine density and found that the group receiving the AAV-Aiolos presented a reduction of spine density compared with the group receiving the AAV-mCherry (Figure 7B). To investigate whether the observed reduction in spine density could be associated with complement-mediated elimination, we next evaluated iC3b abundance and deposition of iC3b within microglia. We found a significant increase in both, total iC3b number of spots and iC3b-positive puncta colocalizing with Iba1-positive microglia in AAV-Aiolos mice compared with controls (Figure 7C-D), consistent with an increased complement-associated microglial engagement. To investigate whether the observed increase in microglia-associated iC3b was related to synaptic loss, we next quantified PSD95 abundance and colocalization of PSD95-positive puncta with Iba1-positive microglia related to total PSD95 levels. We observed a significant reduction of PSD95 number of spots whereas colocalization of PSD95-positive puncta within Iba1-positive microglia was increased in AAV-Aiolos mice compared with controls (Figure 7E-F), suggesting increased microglia-associated engulfment of synaptic material.

**Figure 7.**
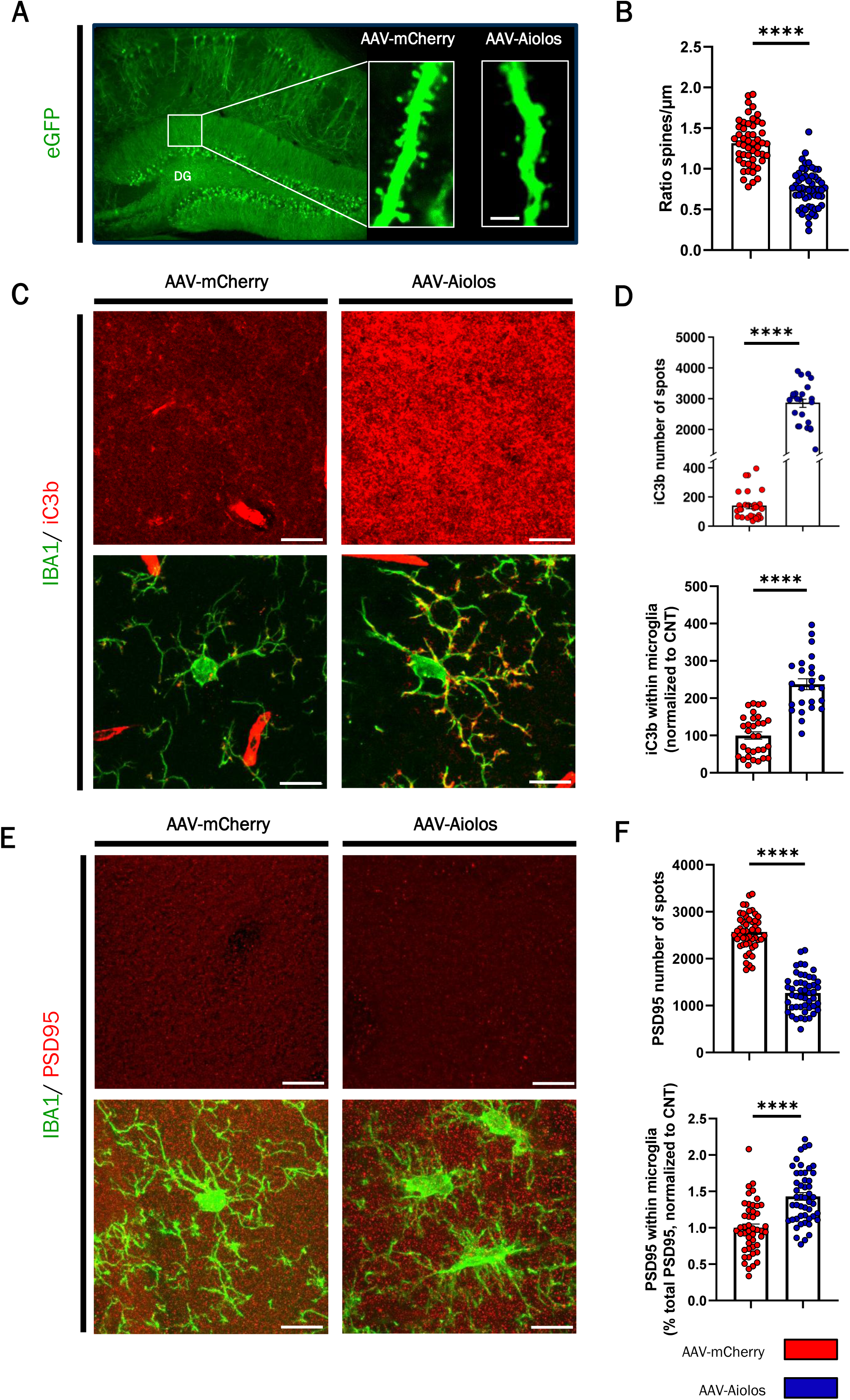
Hippocampal Aiolos overexpression reduces dendritic spine density and increases microglial colocalization of iC3b and PSD95. (A) Representative EGFP fluorescence image of the hippocampus in Thy1-EGFP mice, with higher magnification views of dendritic segments of the dentate gyrus (DG) from mice receiving hippocampal injections of AAV-mCHERRY or AAV-Aiolos. Scale bar, 5 µm. (B) Dendritic spine density in the DG, expressed as the number of spines per µm of dendrite. (C) Representative fluorescence images of iC3b (red) and Iba1-positive microglia (green) in the DG. Scale bars, 10 µm. (D) Quantification of total iC3b-positive puncta (top) and iC3b puncta colocalizing with Iba1, normalized to the AAV-mCHERRY control group (bottom). (E) Representative fluorescence images of PSD95-positive puncta (red, top) and merged images showing PSD95 and Iba1-positive microglia (green, bottom). Scale bars, 10 µm. (F) Quantification of total PSD95-positive puncta (top) and the fraction of total PSD95-positive puncta colocalizing with Iba1, normalized to the AAV-mCHERRY control mean (bottom). Data are presented as mean ± SEM. Each dot represents one dendrite in B or one imaging field acquired as a z-stack in D and F. Five mice were analyzed per group, with at least eight dendrites per mouse in B and at least eight imaging fields per mouse in D and F. Groups were compared using unpaired *t*-tests (B and F) or the Mann–Whitney *U* test (D). **** *p* < 0.0001.

## DISCUSSION

In the present work, we characterized Aiolos expression within the mouse brain across development. We provided in vivo evidence of the presence of endogenous Aiolos protein in the adult mouse brain and found that, within the hippocampus, it localized predominantly in principal neurons. We also found that Aiolos is upregulated in peripheral immune cells and in the hippocampus of patients with schizophrenia but not in patients with major depression and that those dysregulations can be associated with the expression of positive, negative, and cognitive schizophrenia-like phenotypes in mice. Finally, we showed that the overexpression of Aiolos in the hippocampus led to complement upregulation and complement-associated synaptic structural alterations.

Our characterization of Aiolos shows that its expression tends to be higher during perinatal development in many brain regions, resembling the developmental expression patterns described for Ikaros and Helios, other members of Ikaros family. However, Aiolos expression persists in neurons into adulthood, whereas neuronal expression of Ikaros and Helios declines after perinatal development [7], [30], this shows that their expression patterns do not always overlap and suggests that individual Ikaros family members may differently contribute to brain development, maturation and function. Ikaros has also been detected in the adult brain [8], but its restricted expression to microglia remains consistent with its established roles in immune cells. In contrast, the presence of Aiolos in neurons points out that its functions extend beyond the immune system, and in turn suggests distinct roles for the Ikaros family members in different brain cell types. This diversity of functions is further supported by our detection of Aiolos in some astrocytes, although its role in this cell population remains to be addressed in future studies.

Beyond its expression in the healthy brain, we identified Aiolos dysregulations in both peripheral immune cells and the hippocampus of patients with schizophrenia, consistent with the previous reports involving other Ikaros family members in this disorder [9], [10].Our animal models assessing the impact of Aiolos dysregulations shed light on the possibility that these distinct alterations may differentially contribute to behavioral phenotypes relevant to different domains of schizophrenia symptomatology. On the one hand, intracerebroventricular infusion of PBMC-derived secretome from schizophrenia patients selected with elevated Aiolos expression induced the expression of a negative schizophrenia-like behavioral phenotype in WT mice. On the other hand, hippocampal Aiolos overexpression in WT mice generated the expression of positive and cognitive schizophrenia-like behavioral phenotypes. Altogether these results suggest that the immune dysregulation of Aiolos may contribute to the appearance of negative symptoms, while the hippocampal dysregulation of Aiolos preferentially contributes to the appearance of positive and cognitive symptoms. This interpretation is consistent with clinical findings linking peripheral immune alterations, including elevated cytokine levels and splenomegaly, to more severe negative symptoms in patients with schizophrenia [39], [40], [41]. Furthermore, clinical findings have linked altered hippocampal activity to more severe positive symptoms and poorer memory performance in patients with schizophrenia [42]. Consistently, experimental evidence in mice also shows that altering hippocampal activity state can induce positive and cognitive schizophrenia-like phenotypes without social interaction deficits associated with negative symptoms [43].

Given the stronger behavioral phenotype observed in the hippocampal Aiolos overexpression model, we focused our subsequent investigation on the potential underlying molecular alterations. The opposing changes in C1qa and C1qb abundance observed in the hippocampal proteomic profiles of Aiolos KO and Aiolos overexpression models, together with the upregulation of additional complement components, including C1qc and C4b, in the Aiolos overexpression model (Figure6 A-B) suggest that Aiolos regulates the abundance of key complement components. Previous studies have reported similar associations; such as the case of C/EBPβ in astrocytes and DLK in neurons [44], [45]. Indeed, our subsequent in vivo histological analyses showed that Aiolos overexpression induced a marked increase in iC3b abundance, accompanied by loss of dendritic spines and synaptic material, suggesting that Aiolos may modulate complement activity and thereby contribute to these synaptic structural alterations. One possibility is that Aiolos positively modulates neuronal complement expression. Although neurons are not the main source of complement proteins in the brain [46], [47], a previous report has shown that neuronal complement expression can be upregulated by other regulatory proteins [45]. In line with this possibility, C1QC is listed as a predicted Aiolos target in JASPAR-derived motif analyses available through the Harmonizome platform [48], [49], suggesting a potential direct transcriptional link, although this prediction requires experimental validation. Alternatively, Aiolos may indirectly modulate complement by altering neuron–glia signaling, potentially promoting complement production by surrounding glial cells, as reported for neuronal DLK [45]. It could also influence the expression of neuronal complement inhibitors, such as CSMD1 or NPTX2, which limit complement deposition or activity [50], [51]. Such mechanisms could contribute to the increased iC3b abundance and associated synaptic alterations without requiring increased complement synthesis within neurons themselves.

The complement and synaptic structural alterations observed in our Aiolos overexpression model are consistent with the proposed involvement of complement proteins in dysregulated synaptic elimination in schizophrenia [52], [53], [54]. In this context, our findings suggest that Aiolos may regulate complement pathways involved in synaptic elimination, potentially linking its dysregulation to the synaptic alterations associated with schizophrenia. Beyond this interpretation, more speculative possibilities could include the idea that Aiolos participates in synaptic pruning processes. Synaptic pruning contributes to the refinement of neural circuits [55], [56], and dysregulation of this process has been proposed in the context of schizophrenia [57], [58]. Against this background, the higher perinatal expression of Aiolos across several brain regions, together with our findings linking its overexpression to complement-associated loss of synaptic material, raises the possibility that Aiolos may contribute to developmental synaptic pruning. A related possibility is that Aiolos dysregulation also influences the altered synaptic pruning processes proposed in schizophrenia. These possibilities remain speculative but provide directions for future research into the role of Aiolos in physiological and schizophrenia-associated processes.

In conclusion, the present study expands the understanding of Aiolos beyond its established immune roles, by identifying a regulatory role in neurons. Furthermore, our results demonstrate Aiolos dysregulation in schizophrenia and support its implication in the disorder’s pathophysiology, potentially through complement-associated structural synaptic alterations as one of the underlying mechanisms. More broadly, our results highlight the value of neuroimmune perspectives for understanding how peripheral immune and brain-specific alterations may differently contribute to schizophrenia phenotypes expression.

## DATA AVAILABILITY

The data supporting the findings of this study are available from the corresponding authors upon reasonable request.

## ACKNOWLEDGEMENTS

We thank Maria Calvo from the Advanced Microscopy service and Lara Sedó from the Unitat d’Experimentació Animal (CCiT, UB) for their help with confocal microscopy and care and managing the mouse colonies.

## AI DISCLOSURE

During the preparation of this manuscript, the authors used ChatGPT (OpenAI) to assist with language editing. The authors reviewed and edited the text and took full responsibility for the content of the manuscript.

## AUTHOR CONTRIBUTIONS

All authors have approved the manuscript and agree with this submission. Authors contribution: Maoyu Wang, Marcos Galán, Nerea Gómez-Rivada: Writing – review & editing, Writing – original draft, Visualization, Validation, Methodology, Investigation, Formal analysis, Data curation, Conceptualization. Laura López-Molina, Miguel Montero-Escobedo, Pol García-Segura: Methodology, Investigation, Formal analysis. Myriam Cavero, Carla Hernandez, Miqueu Alfonso: Methodology and Investigation. Cristina Malagelada, Jordi Alberch, Kentaro Yomogida, J Javier Meana, Luis F Callado: Resources, Supervision. Belén Arranz, Marina López-Solà: Writing – review & editing, Supervision, Resources, Conceptualization. Iván Ballasch and Albert Giralt: Writing – review & editing, Visualization, Validation, Supervision, Resources, Project administration, Funding acquisition, Conceptualization. All authors have read and approved the article.

## SUPPLEMENTARY TABLES

**Supplementary table 1.** Demographics and illness details for human postmortem hippocampal samples from patients with major depressive disorder and controls.

| Case ID | Diagnostic | Sex | Cause of death | Age | PMI (h) | Medication |
|---|---|---|---|---|---|---|
| 83724 | Control | F | CRF | 50 | 10 | n.a. |
| 176724 | Control | F | Train accident | 36 | 20 | n.a. |
| 51318 | Control | F | Traffic Crash | 71 | 22 | n.a. |
| 61524 | Control | F | CRF | 45 | 12 | n.a. |
| 67551 | Control | F | Hemorrhage | 63 | 10 | n.a. |
| 129945 | Control | F | Traffic Crash | 43 | 3 | n.a. |
| 146709 | Control | F | Accident | 80 | 16 | n.a. |
| 105018 | MDD | F | Suicide | 36 | 32 | Venlafaxine<br>Paroxetine<br>BZD |
| 82233 | MDD | F | Suicide | 70 | 7 | Mirtazapine<br>Sertraline<br>BZD |
| 37221 | MDD | F | CRF | 43 | 24 | Venlafaxine |
| 58221 | MDD | F | Suicide | 49 | 18 | Fluoxetine |
| 121233 | MDD | F | Suicide | 45 | 21 | BDZ |
| 188760 | MDD | F | CFR | 79 | 14 | Amitriptiline |
| 16260 | MDD | F | Suicide | 55 | 24 | n.a. |

**Age (years)** Control (n = 7): 55.43 ± 16.20 \| MDD (n = 11): 53.55 ± 16.10 \| p = 0.8123
**PMI (h)** Control (n = 7): 13.29 ± 6.55 \| MDD (n = 11): 19.82 ± 10.06 \| p = 0.1488
*Values are mean ± SD. P values are from two-sided unpaired Student’s t-tests.*
Abbreviations: CRF: cardiorespiratory failure. AMI: Acute Myocardial Infarction. MDD: Major depressive disorder. BZD: Benzodiazepine. PMI: Post-mortem interval, M: Male, F: Female, (-): Negative results. Toxicology was performed postmortem.

**Supplementary Table 2.** Demographics and illness details for human postmortem hippocampal samples from patients with schizophrenia and controls.

| Case ID | Diagnosis | Sex | Age | PMI (h) | PANSS+ | PANSS- | PANSS G | Medication |
|---|---|---|---|---|---|---|---|---|
| CTR 1 | Control | M | 64 | 3 | n.a. | n.a. | n.a. | n.a. |
| CTR 2 | Control | M | 39 | 3 | n.a. | n.a. | n.a. | n.a. |
| CTR 3 | Control | F | 81 | 23 | n.a. | n.a. | n.a. | n.a. |
| CTR 4 | Control | M | 31 | 17 | n.a. | n.a. | n.a. | n.a. |
| CTR 5 | Control | M | 86 | 7 | n.a. | n.a. | n.a. | n.a. |
| CTR 6 | Control | F | 90 | 13 | n.a. | n.a. | n.a. | n.a. |
| CTR 7 | Control | M | 58 | 5 | n.a. | n.a. | n.a. | n.a. |
| CTR 8 | Control | F | 88 | 24 | n.a. | n.a. | n.a. | n.a. |
| CTR 9 | Control | M | 78 | 5 | n.a. | n.a. | n.a. | n.a. |
| CTR 10 | Control | F | 83 | 7 | n.a. | n.a. | n.a. | n.a. |
| CTR 11 | Control | F | 97 | 7 | n.a. | n.a. | n.a. | n.a. |
| CTR 12 | Control | M | 93 | 5 | n.a. | n.a. | n.a. | n.a. |
| CTR 13 | Control | F | 83 | 7 | n.a. | n.a. | n.a. | n.a. |
| CTR 14 | Control | M | 86 | 7 | n.a. | n.a. | n.a. | n.a. |
| SCH 1 | SCZ | M | 34 | 2 | --- | --- | --- | No treatment |
| SCH 2 | SCZ | M | 76 | 7 | 12 | 18 | 37 | Quetiapine |
| SCH 3 | SCZ | M | 80 | 4,5 | 12 | 28 | 34 | Olanzapine |
| SCH 4 | SCZ | M | 86 | 1 | --- | --- | --- | Levomepromacine |
| SCH 5 | SCZ | M | 77 | 2 | 11 | 23 | 50 | Risperidone and Quetiapine |
| SCH 6 | SCZ | M | 82 | 6 | --- | --- | --- | Risperidone |
| SCH 7 | SCZ | M | 79 | 7 | --- | --- | --- | Haloperidol |
| SCH 8 | SCZ | M | 87 | 8,5 | 16 | 41 | 59 | n.a. |
| SCH 9 | SCZ | M | 83 | 4 | 11 | 29 | 47 | Clorpromazine and Haloperidol |
| SCH 10 | SCZ | M | 90 | 3 | --- | --- | --- | No treatment |
| SCH 11 | SCZ | M | 86 | 6 | --- | --- | --- | n.a. |
| SCH 12 | SCZ | M | 84 | 3 | 29 | 47 | 62 | Amisulpride |
| SCH 13 | SCZ | M | 82 | 7 | --- | --- | --- | n.a. |
| SCH 14 | SCZ | M | 91 | 3 | 35 | 42 | 65 | No treatment |

**Age (years)** Control (n = 14): 75.50 ± 20.11 \| SCZ (n = 14): 79.79 ± 13.91 \| p = 0.5177
**PMI (h)** Control (n = 14): 9.50 ± 7.00 \| SCZ (n = 14): 4.57 ± 2.34 \| p = 0.0191
*Values are mean ± SD. P values are from two-sided unpaired Student’s t-tests.*
Abbreviations: *post-mortem* interval (PMI), Positive and Negative Syndrome Scale (PANSS). Schizophrenia (SCZ)

**Supplementary Table 3.** Demographics and illness details from schizophrenia patients and controls of whom we obtained the peripheral blood mononuclear cells (PBMCs)

| Case ID | Diagnosis | Sex | Age | PANSS+ | PANSS- | PANS G | Medication |
|---|---|---|---|---|---|---|---|
| CTR 1 | Control | F | 27 | n.a. | n.a. | n.a. | n.a. |
| CTR 2 | Control | F | 37 | n.a. | n.a. | n.a. | n.a. |
| CTR 3 | Control | M | 38 | n.a. | n.a. | n.a. | n.a. |
| CTR 4 | Control | M | 37 | n.a. | n.a. | n.a. | n.a. |
| CTR 4 | Control | M | 42 | n.a. | n.a. | n.a. | n.a. |
| CTR 5 | Control | F | 31 | n.a. | n.a. | n.a. | n.a. |
| CTR 6 | Control | F | 42 | n.a. | n.a. | n.a. | n.a. |
| CTR 7 | Control | M | 24 | n.a. | n.a. | n.a. | n.a. |
| CTR 8 | Control | M | 26 | n.a. | n.a. | n.a. | n.a. |
| CTR 9 | Control | F | 26 | n.a. | n.a. | n.a. | n.a. |
| CTR 10 | Control | M | 26 | n.a. | n.a. | n.a. | n.a. |
| CTR 11 | Control | F | 24 | n.a. | n.a. | n.a. | n.a. |
| CTR 12 | Control | M | 60 | n.a. | n.a. | n.a. | n.a. |
| CTR 13 | Control | F | 55 | n.a. | n.a. | n.a. | n.a. |
| CTR 14 | Control | M | 53 | n.a. | n.a. | n.a. | n.a. |
| CTR 15 | Control | M | 52 | n.a. | n.a. | n.a. | n.a. |
| SCH 1 | SCZ | M | 35 | 12 | 19 | 31 | Paliperidone |
| SCH 2 | SCZ | M | 43 | 14 | 22 | 29 | Clozapine |
| SCH 3 | SCZ | F | 43 | 7 | 18 | 26 | Clozapine |
| SCH 4 | SCZ | M | 27 | 9 | 24 | 36 | Clozapine |
| SCH 5 | SCZ | M | 25 | 15 | 18 | 26 | Clozapine |
| SCH 6 | SCZ | M | 40 | 21 | 18 | 36 | Olanzapine |
| SCH 7 | SCZ | M | 29 | 11 | 27 | 40 | Aripiprazol |
| SCH 8 | SCZ | F | 22 | 10 | 33 | 34 | Aripiprazol |
| SCH 9 | SCZ | F | 58 | - | - | - | Paliperidone |
| SCH 10 | SCZ | F | 48 | 17 | 26 | 39 | Clozapine |
| SCH 11 | SCZ | F | 37 | 14 | 28 | 34 | Clozapine |
| SCH 12 | SCZ | F | 56 | 9 | 29 | 33 | Clozapine |
| SCH 13 | SCZ | M | 46 | 14 | 25 | 29 | Clozapine |
| SCH 14 | SCZ | F | 54 | 10 | 21 | 27 | Clozapine |
| SCH 15 | SCZ | F | 48 | 19 | 25 | 32 | Clozapine |
| SCH 16 | SCZ | M | 45 | 7 | 11 | 20 | Clozapine |
| SCH 17 | SCZ | F | 49 | 18 | 36 | 49 | Clozapine |
| SCH 18 | SCZ | M | 32 | 7 | 19 | 25 | Clozapine |
| SCH 19 | SCZ | M | 39 | - | - | - | Paliperidone |
| SCH 20 | SCZ | F | 48 | - | - | - | Olanzapine |
| SCH 21 | SCZ | M | 33 | 10 | 16 | 27 | Clozapine |
| SCH 22 | SCZ | F | 37 | 11 | 20 | 27 | Clozapine |
| SCH 23 | SCZ | F | 28 | 12 | 14 | 27 | Paliperidone |
| SCH 24 | SCZ | M | 50 | 18 | 19 | 40 | Paliperidone |
| SCH 25 | SCZ | F | 19 | 9 | 29 | 22 | Clozapine |
| SCH 26 | SCZ | F | 37 | 32 | 11 | 49 | Olanzapine |
| SCH 27 | SCZ | F | 51 | 12 | 34 | 38 | Paliperidone |
| SCH 28 | SCZ | M | 25 | 14 | 14 | 28 | Clozapine |
| SCH 29 | SCZ | M | 18 | - | - | - | Olanzapine |
| SCH 30 | SCZ | M | 27 | 7 | 11 | 28 | - |
| SCH 31 | SCZ | F | 35 | 7 | 14 | 24 | Aripiprazol |
| SCH 32 | SCZ | F | 40 | 7 | 16 | 16 | Olanzapine |
| SCH 33 | SCZ | F | 54 | 14 | 29 | 35 | Aripiprazol |
| SCH 34 | SCZ | M | 23 | 13 | 13 | 19 | Paliperidone |
| SCH 35 | SCZ | M | 57 | 7 | 27 | 19 | Risperidone |

**Age (years)** Control (n = 16): 37.50 ± 12.16 \| SCZ (n = 35): 38.80 ± 11.59 \| p = 0.7159
*Values are mean ± SD. P values are from two-sided unpaired Student’s t-tests.*
Abbreviations: Positive, Negative and General Syndrome Scale (PANSS). Schizophrenia (SCZ)

**Supplementary Table 4.** Demographics and illness details from major depressive disorder patients and controls of whom we obtained the peripheral blood mononuclear cells (PBMCs)

| Case ID | Diagnostic | Sex | Age | Years of schooling |
|---|---|---|---|---|
| 10 | Control | F | 50 | 15 |
| 14 | Control | F | 53 | 18 |
| 17 | Control | F | 38 | 28 |
| 25 | Control | F | 53 | 12 |
| 39 | Control | F | 54 | 14 |
| 42 | Control | F | 48 | 30 |
| 45 | Control | F | 52 | 22 |
| 49 | Control | F | 59 | 17 |
| 53 | Control | F | 62 | 15 |
| 55 | Control | F | 51 | 25 |
| 56 | Control | F | 47 | 19 |
| 61 | Control | F | 43 | 20 |
| 64 | Control | F | 56 | 17 |
| 66 | Control | F | 54 |  |
| 68 | Control | F | 50 | 14 |
| 69 | Control | F | 57 | 20 |
| 70 | Control | F | 53 | 18 |
| 73 | Control | F | 43 | 25 |
| 91 | Control | F | 43 | 24 |
| 98 | Control | F | 61 | 11 |
| 2 | MDD | F | 46 | 18 |
| 32 | MDD | F | 59 | 13 |
| 47 | MDD | F | 50 | 10 |
| 54 | MDD | F | 43 | 26 |
| 57 | MDD | F | 46 | 20 |
| 60 | MDD | F | 57 | 22 |
| 72 | MDD | F | 55 | 24 |
| 80 | MDD | F | 31 | 26 |
| 81 | MDD | F | 63 | 17 |
| 83 | MDD | F | 31 | 19 |
| 84 | MDD | F | 65 | 21 |
| 92 | MDD | F | 55 | 15 |
| 93 | MDD | F | 58 | 21 |
| 99 | MDD | F | 33 | 20 |

**Age (years)** Control (n = 20): 51.35 ± 6.34 \| MDD (n = 14): 49.43 ± 11.52 \| p = 0.5362
*Values are mean ± SD. P values are from two-sided unpaired Student’s t-tests.*

**Supplementary Table 5.**
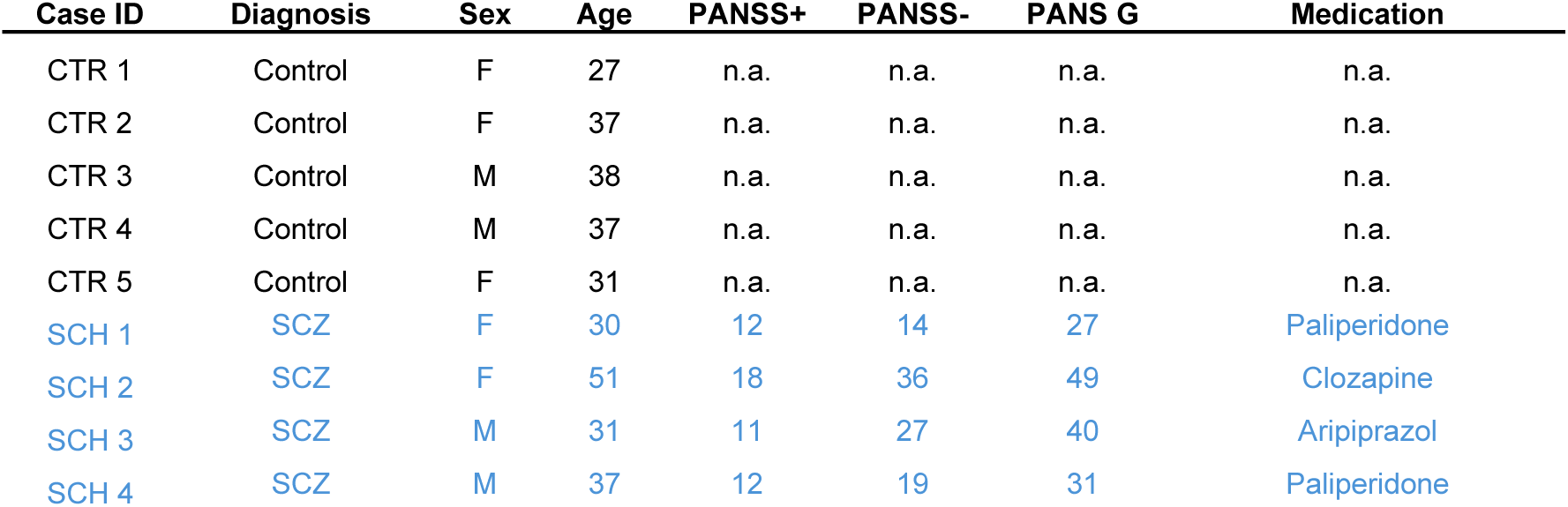
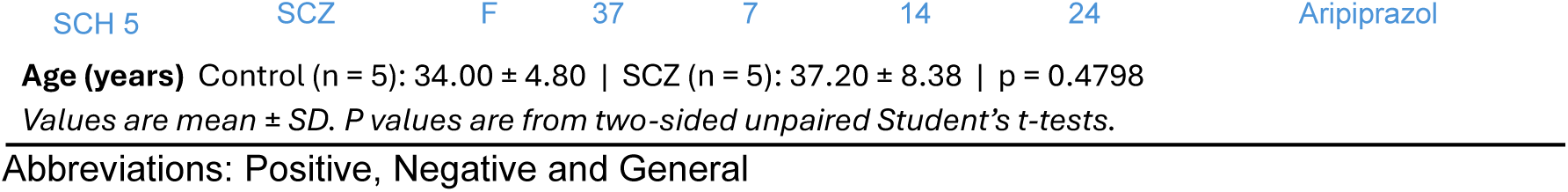
Demographics and illness details from schizophrenia patients and controls of whom we obtained the peripheral blood mononuclear cells (PBMCs) and the secretome. We specifically selected patients with an up-regulation of Aiolos (> 40%) in PBMCs compared to the control subjects.

| Case ID | Diagnosis | Sex | Age | PANSS+ | PANSS- | PANS G | Medication |
|---|---|---|---|---|---|---|---|
| CTR 1 | Control | F | 27 | n.a. | n.a. | n.a. | n.a. |
| CTR 2 | Control | F | 37 | n.a. | n.a. | n.a. | n.a. |
| CTR 3 | Control | M | 38 | n.a. | n.a. | n.a. | n.a. |
| CTR 4 | Control | M | 37 | n.a. | n.a. | n.a. | n.a. |
| CTR 5 | Control | F | 31 | n.a. | n.a. | n.a. | n.a. |
| SCH 1 | SCZ | F | 30 | 12 | 14 | 27 | Paliperidone |
| SCH 2 | SCZ | F | 51 | 18 | 36 | 49 | Clozapine |
| SCH 3 | SCZ | M | 31 | 11 | 27 | 40 | Aripiprazol |
| SCH 4 | SCZ | M | 37 | 12 | 19 | 31 | Paliperidone |
| SCH 5 | SCZ | F | 37 | 7 | 14 | 24 | Aripiprazol |

**Age (years)** Control (n = 5): 34.00 ± 4.80 | SCZ (n = 5): 37.20 ± 8.38 | p = 0.4798
*Values are mean ± SD. P values are from two-sided unpaired Student’s t-tests.*
Abbreviations: Positive, Negative and General

